# A Longitudinal Study of Dominant *E. coli* Lineages and Antimicrobial Resistance in the Gut of Children Living in an Upper Middle-income Country

**DOI:** 10.1101/2022.01.11.475974

**Authors:** Diana Calderón, Paúl A. Cárdenas, Belen Prado, Jay P. Graham, Gabriel Trueba

## Abstract

The gastrointestinal tract constitutes a complex and diverse ecosystem. *Escherichia coli* is one of the most frequently studied and characterized species in the gut ecosystem, nevertheless, there has been little research to determine their diversity and population dynamics in the intestines of children over time. In this prospective study, a fresh fecal sample was obtained from children longitudinally over one year (30 fecal samples at sampling period 1 and 22 fecal samples at sampling periods 2 and 3). From each stool sample, five *E. coli* colonies were randomly selected (n = 405 *E. coli* isolates total) in order to characterize the genotype and phenotypic antimicrobial resistance patterns. We found that all numerically dominant *E. coli* lineages in children’s intestines were transient colonizers, and antimicrobial resistance phenotypes of these strains varied significantly over time without any apparent selective force. Whole-genome sequencing of 3 isolates belonging to ST131 found in one child during the sampling period I and II indicated that isolates were three different ST 131 clones that carried extended-spectrum β-lactamase (ESBL) genes.

**IMPORTANCE:** The length of residency and numeric dominance of antimicrobial-resistant *E. coli* may affect the extent to which an isolate contributes to the dissemination of antimicrobial resistance in a community. We studied the persistence of numerically dominant and antimicrobial-resistant lineages of *E. coli* in the human intestine. We found that *E. coli* lineages in the children's gut change considerably and rapidly over time. This study suggests that some phenotypic resistance patterns may result from the random distribution of genes in *E. coli* populations over time and may not be associated with differential exposure to antimicrobials.

## INTRODUCTION

*Escherichia coli* is a minor component of the intestinal microbiota of warm-blooded animals [1, and 2]. Within the human gut and other warm-blooded animals, there may be more than 16 different lineages of *E. coli* at any time [3]; each of these lineages is present in different abundances and remains in the intestine for different periods. Intestinal *E. coli* could be classified as follow: resident when the lineage remains in the intestine for months and even years [4], transient when the lineage remains for days or weeks [4], dominant when the isolates make up a large proportion (> 50%) [5] of the *E. coli* cells in the intestine, and minority when the proportion is smaller (< 10%) [3].

The genetic dynamics of the *E. coli* population in the intestine have received little attention, even though some are a source of problematic infections and antimicrobial resistance genes (ARGs) [6, 7, 8]. The high abundance and the persistence of certain *E. coli* pathogenic and antimicrobial-resistant lineages in the intestinal tract have been suggested to be a critical risk factor for disease and disease treatment [ 8, 9, 10]. Furthermore, these lineages could affect the dissemination of ARGs since *E. coli* is a species adept at horizontal gene transfer [1, 2, 11] and is likely the intestinal species that can be exchanged the most between different hosts [12].

Here we screened the dominant *E. coli* strains obtained from the feces of children less than five years of age and we also analyzed the turnover of these strains in three sampling periods (SP I, II, and III) during one year; sampling periods were approximately 3 months apart. Dominant *E. coli* strains were defined as those colonies that grew on MacConkey agar at the highest proportion [5].

## RESULTS

We analyzed 405 *E. coli* isolates from 82 fecal samples: 30 children were sampled in SP I and SP II and 22 were sampled SP I through SP III. Clonal relationships were first assessed by sequencing the *fum*C alleles [14]: those isolates that carried a different *fumC* allele were classified as non-clonal. The isolates that were obtained from the same child in different SPs and showed the same *fumC* allele were subjected to MLST analysis to determine the isolate's sequence type (ST). Isolates that showed a different ST number were also classified as non-clonal, while the isolates with the same ST number were analyzed using whole-genome sequencing (WGS) to confirm clonality. All isolates were subjected to antimicrobial susceptibility test following CLSI guidelines [13].

Among the 405 isolates, we found a total of 40 different *fumC* alleles where the most prevalent were *fumC* 11 (n=108, 26.7%), *fumC* 35 (n=40, 9.9%) and *fumC* 4 (n=38, 9.4%). Among the dominant *E. coli* isolates randomly selected from each fecal sample, 47 of 82 fecal samples had the same *fumC* allele in all five colonies; 12 of the 47 fecal samples had isolates with the *fumC* 11 allele (14.6%), followed by 5 fecal samples that had isolates with the *fumC* 35 allele and 5 with the *fumC* 4 allele (6.1% respectively).

We identified 14 cases (samples obtained from 12 children) in which we obtained isolates with the same *fumC* allele in different SPs from the same individual, however when we carried out MLST, there were 4 cases where the isolates were the same ST (ST34 was recovered twice, 11 months apart in child 26; ST10 was recovered twice in children 20 and 30) at 7 and 5 months apart respectively, and ST131 was recovered twice in child 23, 4 months apart) (Fig. 1). Whole-genome sequencing showed that none of the isolates belonging to the same ST were clonal. In SPI (child 23) we found E. coli isolates carrying *fumC* 40 in SPI and SPII. All *fumC* 40 isolates were resistant to imipenem except for 1 isolate (collected in SPI) that was sensitive to this antibiotic. MLST and WGS from 2 *fumC* 40 isolates (from SPI and SPII) and the *fumC* 40 isolate sensitive to imipenen showed that the 3 isolates belonged to ST131 however none of the isolates was clonal ( we found 589 SNPs among the ST131 isolates) (Figure 2).

**FIGURE 1.**
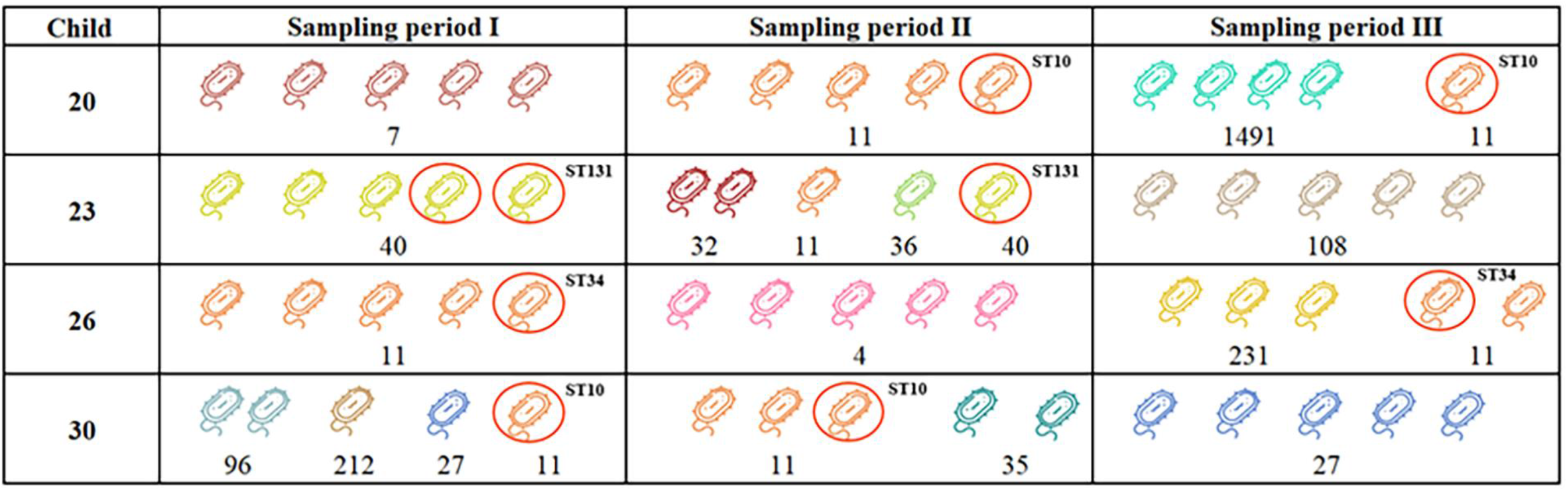
Numerically dominant *E. coli* isolates showing evidence of permandence in children intestines for 3 months. The figure showed 5 isolates obtained from 4 children during 3 periods (aproximatelly 3 months apart). The numbers indicate different alleles of the *fumC* gene. The isolates showing same *fumC* allele during 3 periods were subjected to MLST and represented by sequence types (ST)

**FIGURE 2:**
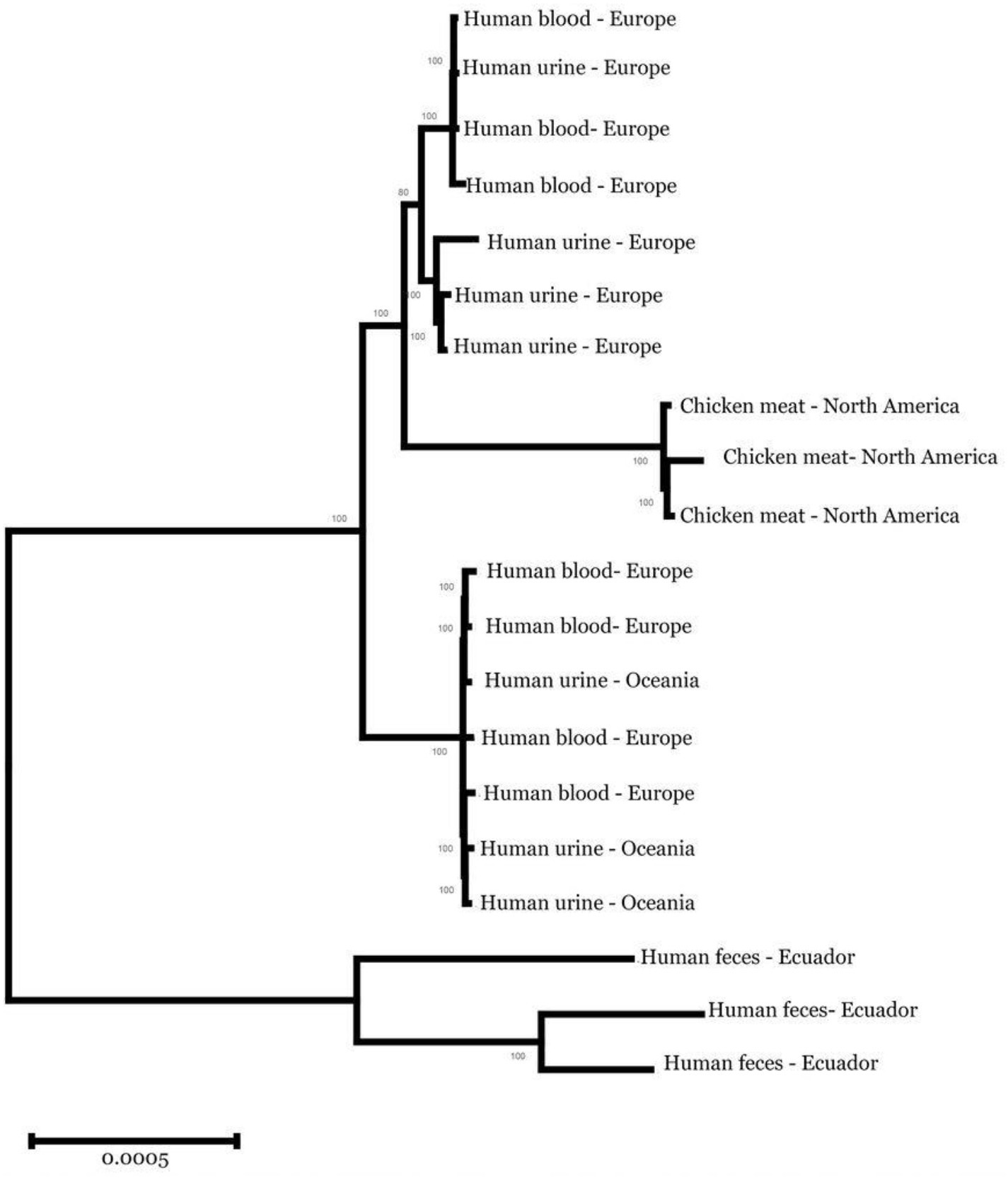
Maximun Likelihood tree of whole genome sequences of *E. coli* isolates belonging to ST131. The analysis shows our sequences from Ecuador compared with sequences of *E. coli* from different sources in North America, Europe, and Oceania.

Among the 405 isolates recovered, 122 (30.1%) were susceptible to all the antimicrobials tested. The remaining fell into one of the 75 unique antibiotic resistance profiles: 50 isolates (12.3%) were resistant to only one antibiotic, 37 isolates (9.1%) were resistant to two antibiotics, and 196 (48.4%) were resistant to three or more antimicrobials. Phenotypic resistance to the combination of ampicillin (AM), trimethoprim-sulfamethoxazole (SXT), and tetracycline (TE) was the most common profile (n=36, 8.9%), followed by tetracycline (TE) resistance (n=21, 5.2%), and by the combination of ampicillin (AM), trimethoprim-sulfamethoxazole (SXT), and amoxicillin-clavulanic acid (AMC) resistance (n=15, 3.7%) (Table 1).

**TABLE 1.** Prevalence of phenotypical resistance of *Escherichia coli* numerically dominant isolates obtained from 22 children during SP I, SP II and SP III. Bold numbers indicate statistic significance

| Antimicrobials | SPI | SPII | SPIII | X<sup>2</sup> test | p value |
| --- | --- | --- | --- | --- | --- |
|  | Oct.-Dec. 2018 | Jan.-May 2019 | Jul.-Dec 2019 |  |  |
| Ampicillin (AM) | 64 (58.2 %) | 60 (54.5 %) | 46 (41.8 %) | **6.502** | **0.039** |
| Chloramphenicol (C) | 5 (4.5 %) | 12 (10.9 %) | 9 (8.2 %) | 3.090 | 0.213 |
| Imipenem (IPM) | 12 (10.9 %) | 1 (0.9 %) | 7 (6.4 %) | **9.687** | **0.008** |
| Trim-sulfamet (SXT) | 58 (52.7 %) | 58 (52.7 %) | 50 (45.4 %) | 1.552 | 0.460 |
| Gentamicin (GM) | 0 (0 %) | 6 (5.4 %) | 0 (0 %) |  |  |
| Ceftazidime (CAZ) | 2 (1.8 %) | 14 (12.7 %) | 10 (9.1 %) | **9.352** | **0.009** |
| Cefepime (FEP) | 13 (11.8 %) | 15 (13.6 %) | 9 (8.2 %) | 1.705 | 0.426 |
| Cefazolin (CZ) | 18 (16.4 %) | 14 (12.7 %) | 15 (13.6 %) | 0.645 | 0.724 |
| Ciprofloxacin (CIP) | 22 (20 %) | 21 (19.1 %) | 9 (8.2 %) | **7.168** | **0.028** |
| Cefotaxime (CTX) | 14 (12.7 %) | 18 (16.4 %) | 14 (12.7 %) | 0.808 | 0.668 |
| Amoxicillin + Clav. Ac.(AMC) | 35 (31.8 %) | 28 (25.4 %) | 15 (13.6 %) | **10.376** | **0.006** |
| Tetracycline (TE) | 55 (50 %) | 52 (47.3 %) | 50 (45.4 %) | 0.462 | 0.794 |

(P value < 0.05)

There were statistically significant differences in the resistance to chloramphenicol (C) between SP I and SP II, with a higher resistance in SP I (Table 1). Similarly, statistically significant differences in the resistance to ceftazidime (CAZ) were found, with a higher resistance in SP II versus SP III (Table 1 and Table S3).

## DISCUSSION

In this one-year prospective study, we followed 30 children for three sampling periods to screen the predominant *E. coli* strains present in fresh fecal samples. We found that all numerically dominant *E. coli* strains were transient colonizers, which suggests a high turnover rate of this species’ lineages in the gut of children (Figure 1). This observation agrees with previous reports showing a high diversity and high turnover rate of *E. coli* lineages in the human intestines [15, 16, 17, 18]. It remains unknown what causes this high strain turnover; however, bacteriophage infection, bacterial bacteriocins, protozoal predation, immune mechanisms, and diet are probably key factors [1, 15, 19]. We also found that at least three of the five selected colonies in every sample showed the same antimicrobial resistance profile as well as the same *fumC* allele, which suggests that the large proportion of the isolates obtained at a specific time, from one individual, were the same clone; this is consistent with the idea that many of these colonies represent dominant strains [5]. We found 3 different clones of ST131 colonizing one child (Individual #23) collected during 2 different sampling periods; the isolates showed phenotypic resistance to the AM-SXT-FEP-CZ-CIP-CTX-AMC antimicrobials, 2 of them (isolated during 2 different periods) showed resistance to imipenem (IMP). ST 131 is a known human extra-intestinal pathogen, characterized by its virulence [20] and by its resistance to extended-spectrum cephalosporins and fluoroquinolones [20 and 21]. In the same way, it is strongly related to extra-intestinal infections, mainly urinary tract, which has been reported as dominant in human intestines [22], domestic animal intestines, and the environment [20, 21, 22]; our isolates belonged to a different cluster from those associated with humans or domestic animals (Figure 2). It is unclear why this child was repeatedly colonized by ST131, however, the child had a diarrhea episode and was exposed to antibiotics in SP I, and visited a private clinic before SP I and SP III; it is possible that this child acquired these ST131 clones in the first visit to the health care facility. Our results also corroborated the notion that ST131 is a successful epidemic clone [22].

Among the 405 isolates analyzed, 30.1% (122 of 405) were susceptible to all of the antimicrobials tested. Resistance to ampicillin (AM), trimethoprim-sulfamethoxazole (SXT), and tetracycline (TE) were the most prevalent, while gentamycin (GM) resistance was the least common; only six strains showed resistance to GM. We found 24 (5.9%) *E. coli* isolates (including one ST131 isolate) were resistant to IPM and 40 isolates (9.8%) to cefepime (FEP); these antibiotics are used uniquely in humans and the resistances should be associated with hospitals and not community isolates. In the imipenem (IMP) resistant ST131 isolates we found *bla*_OXA-1_, *bla*_OXA-368_, and *bla*_CTX-M15_ genes whereas in the IMP sensitive ST 131 isolate, the *bla*_OXA-368_ was missing; this suggests that the *bla*_OXA-368_ in combination with another ESBL gene or a porin mutation may be responsible for the IMP resistance phenotype [23].

There were significant different resistance to ampicillin (AM), ceftazidime (CAZ), ciprofloxacin (CIP), and amoxicillin + clavulanic acid (AMC) among SP I, SP II and SP III (Table 1). Our household survey that was designed to capture antibiotic use in the household, as well as other risk factors, did not detect any potential cause for this difference in antimicrobial susceptibility. For example, six children (20%) were treated with antimicrobials in SP I, 5 (16.6%) children received antimicrobials in SPII and 5 (22.7%) of the children received antimicrobials in SPIII. Interestingly, one child (individual #23) received antimicrobials repeatedly in SP I and SP III and was found colonized with 3 multi-drug resistant ST131 isolates. All 30 households reported access to uninterrupted potable water, and 28 (93.3%) had functional sewerage systems. During the sampling periods, 8 children (26.7%) changed their dietary habits (stopped breastfeeding) in SP I, 14 (46.7%) reported having diarrhea in the last seven days prior to sampling, and 16 (53.3%). Our findings suggest that *E. coli* antimicrobial resistant-prevalence surveys are potentially not that useful because the phenotypic resistance profiles change rapidly over time, without any identifiable driving factors. These results may be in line with some antimicrobial resistance reports that show similar levels of some antimicrobial resistance in animals treated with antimicrobials and those not given antimicrobials [24].

Understanding the factors involved in lineage turnover is critically important for understanding antimicrobial resistance and virulence gene carriage in commensal strains. Our research shows another level of complexity on the understanding of the gut microbiome, which is important as many strains of the same species carry different metabolic, virulence, or antibiotic resistance genes. Our study reinforces the idea of the constant transition of microbiome members over time [1, 2, 16, 25] and the need for more research in the dynamics of strain populations in the intestine.

## MATERIALS AND METHODS

### Study locations

This one-year longitudinal study was carried out in six semi-rural communities belonging to the parishes of Yaruquí, Pifo, Tumbaco, Checa, Puembo, and Tababela; all of them located near Quito, Ecuador. For household enrollment, inclusion criteria included: (i) households have a child aged six months to four years, (ii) households have a childcare provider who was over eighteen years, and (iii) informed consent was provided by the primary childcare provider. Sixty-one households were enrolled at the beginning of the study but, only 26 finished the full longitudinal study.

### Ethical considerations

The study protocol was approved by the Committee for Protection of Human Subjects (CPHS) and the Office for Protection of Human Subjects (OPHS) at the University of California, Berkeley (Federalwide Assurance # 6252), the Human Research Ethics Committee at the Universidad San Francisco de Quito (no. 2017-178M) and the Ministry of Public Health, Ecuador (MSPCURI000243-3).

### Sample collection

We collected a single fecal sample from each child during three sampling periods (SP): from October to December of 2018 (SP I), from January to May of 2019 (SP II), and from July to December of 2019 (SP III), obtaining a total of 120 stool samples. Each time a sample was collected, the childcare provider completed a survey related to the current family lifestyle and recent exposure-related factors relevant to AMR such as domestic animal, child antimicrobial use, accessibility to water, sanitation, and hygiene (WaSH) conditions (the results of this survey was published previously [7, 28].

### *Escherichia coli* isolation

Each fecal sample was plated on MacConkey agar and incubated at 37 °C for 18 hours. To ensure the selection of the dominant *Escherichia coli* strains, we collected 5 colonies (with same morphology) which provides more than 95% likelihood of obtaining a dominant strain (comprising ≥ 50% of all *E. coli* colonies that grow in a Petri dish from a fecal sample) [26]. Additionally, each colony was transferred to Chromocult^®^ coliform agar for the identification of *Escherichia coli* through its β-D-glucuronidase activity. Those strains were incubated in Brain Heart Infusion (BHI) medium + glycerol (15%) at 37 °C for 18 hours to perform the antimicrobial susceptibility test. After that, the tubes were stored at −80 °C (30).

### Antimicrobial susceptibility test

We used the Kirby Bauer technique (disc diffusion in Muller Hinton agar) to determine the strains antimicrobial susceptibility using the following twelve antimicrobial discs: cefazolin (CZ; 30 μg), ciprofloxacin (CIP; 5 μg), ampicillin (AM; 10 μg), chloramphenicol (C; 30 μg), imipenem (IPM; 10 μg), trimethoprim-sulfamethoxazole (SXT; 1.25/23.75 μg), gentamicin (GM; 10 μg), ceftazidime (CAZ; 30 μg), cefepime (FEP; 30 μg), cefotaxime (CTX; 30 μg), tetracycline (TE; 30 μg) and amoxicillin + clavulanic acid (AMC; 20/10 μg). Resistance or susceptibility was determined according to Clinical and Laboratory Standards Institute (CLSI) guidelines [13].

### DNA extraction

Each isolate was grown on MacConkey agar at 37 °C for 18 hours and 5-6 colonies were placed into Eppendorf^®^ tubes with 500 μl of sterile distilled water and DNA was released by boiling cells suspensions for 1 minute. The quality of the DNA was monitored by gel electrophoresis [7].

### Strains Genotyping

The clonal relationship of the isolates were determined by amplifying and sequencing the *fumC* gene [21] using the Master Mix Go Taq. Those isolates coming from the same individual and sharing the same *fumC* allele were subjected to full multi-locus sequence-typing (MLST). Briefly, PCR conditions were: 180 secs at 95°C, 30 cycles of 30 secs at 94°C, 30 secs at the annealing temperature of each primer (*adk*: 52 °C; *fumC*: 55 °C; *gyrB* and *mdh*: 58 °C; *icd* and *recA*: 54 °C; *purA*: 50 °C) and 60 secs at 72 °C (*fumC* and *icd*) or 45 secs at 72 °C (*adk*, *gyrB*, *mdh*, *purA*, and *recA*), and a final extension of 7 minutes at 72 °C.

### DNA sequencing

All PCR products were sequenced at Macrogen Inc. using the Sanger sequencing method. The sequences were analyzed using the program Geneious Prime 2020 and were screened using the Enterobase database [26].

### Whole Genome Sequencing

Genomic DNA was extracted and purified from 9 isolates (2 from child 20, 26 and 30; and 3 from child 23) using the Wizard^®^ Genomic DNA Purification according to the manufacturer’s instructions. WGS was carried out using the Oxford Nanopore Technology (ONT) Rapid Barcoding Sequencing protocol. Genome assembly was performed using the Flye assembler while the genome annotation was carried out using Prokka. Acquired AMR genes were identified using ABRicate. ARG-ANNOT, Resfinder and CARD databases were used for the identification of resistance genes.

### Phylogenetic analysis

We searched for whole genomes of *E. coli* ST131 from different parts of the world and from isolates obtained from humans and domestic animals. Pan-genome analysis was carried out using Roary and a maximum-likelihood phylogenetic tree with 1,000 bootstrap replicates based on core genomes of isolates was created using RaxML-NG [29]. Sequence data from additional strains were included in this analysis and were obtained from the Enterobase database [26].

### Statistical analysis

Significant differences between phenotypic antimicrobial resistance prevalence of the individuals through time were tested using a chi-square test [27].

## ACKNOWLEDGMENTS

P.C. is funded by NIH FIC D43TW010540 Global Health Equity Scholars. The study was also supported by the National Institutes of Health under Award Number R01AI135118. The content is solely the responsibility of the authors and does not necessarily represent the official views of the National Institutes of Health. The funders had no role in study design, data collection, and interpretation, or the decision to submit the work for publication.

## REFERENCES

1. Tenaillon O, Skurnik D, Picard B, Denamur E. 2010. The population genetics of commensal *Escherichia coli*. Nat Rev Microbiol 8:207–217 doi:101038/nrmicro2298

2. Priya S, Blekhman R. 2019. Population dynamics of the human gut microbiome: change is the only constant. Genome Biol 31:150 doi: 10.1186/s13059-019-1775-3.

3. Schlager TA, Hendley JO, Bell AL, Whittam TS. 2002. Clonal diversity of *Escherichia coli* colonizing stools and urinary tracts of young girls. Infect Immun 70:1225–1229 doi:101128/iai7031225-12292002

4. Caugant, D, Levin B, Selander, R. 1981. Genetic diversity and temporal variation in the *E coli* population of a human host. Genetics 98:467–490

5. Lautenbach E, Bilker W, Tolomeo P, Maslow J. 2008. Impact of diversity of colonizing strains on strategies for sampling *Escherichia coli* from fecal specimens. J Clin Microbiol 46:3094–3096 doi:101128/jcm00945-08

6. Levin B, Lipsitch M, Perrot V, Schrag S, Antia R, Simonsen L, Moore N, Stewart F. 1997. The population genetics of antibiotic resistance. Clin Infec Dis 24:S9–S16

7. Salinas L, Cárdenas P, Johnson T, Vasco K, Graham J, Trueba G. 2019. Diverse commensal *Escherichia coli* and plasmids disseminate antimicrobial resistance genes in domestic animals and children in a semirural community in Ecuador. mSphere 4:e00316–19 doi:101128/msphere00316-19

8. Peña-Gonzalez A, Soto-Girón MJ, Smith S, Sistrunk J, Montero L, Páez M, Ortega E, Hatt JK, Cevallos W, Trueba G, Levy K, Konstantinidis KT. 2019. Metagenomic signatures of gut infections caused by different *Escherichia coli* pathotypes. Appl Environ Microbiol 85:e01820–19

9. Montealegre M, Talavera A, Roy S, Iqbal M, Aminul M, Lanza V, Julian T. 2020. High genomic diversity and heterogenous origins of pathogenic and antibiotic-resistant *Escherichia coli* in household settings represent a challenge to reducing transmission in low-income settings. mSphere 5:e00704–19

10. Davies N, Flasche S, Jit M, Atkins K. 2019. Within-host dynamics shape antibiotic resistance in commensal bacteria. Nat Ecol Evo 3:440–449. doi:101038/s41559-018-0786-x

11. de Been M, Lanza VF, de Toro M, Scharringa J, Dohmen W, Du Y, Hu J, Lei Y, Li N, Tooming-Klunderud A, Heederik DJJ, Fluit AC, Bonten MJM, Willems RJL, de la Cruz F, van Schaik W. 2014. Dissemination of cephalosporin resistance genes between Escherichia coli strains from farm animals and humans by specific plasmid lineages. PLoS Genet 10:e1004776.

12. Moeller AH, Suzuki TA, Phifer-Rixey M, Nachman MW. 2018 Transmission modes of the mammalian gut microbiota. Science. 362:453–457.

13. Clinical and Laboratory Standards Institute Performance Standards for Antimicrobial Susceptibility Testing 2018 28^th^ Ed CLSI supplement M100

14. Barrera S, Cárdenas P, Graham J, Trueba G. 2018. Changes in dominant *Escherichia coli* and antimicrobial resistance after 24 hr in fecal matter. MicrobiologyOpen e00643 doi:101002/mbo3643

15. Martinson J, Pinkham N, Peters G, Cho H, Heng J, Rauch M, Broadaway S, Walk S. 2019. Rethinking gut microbiome residency and the Enterobacteriaceae in healthy human adults. ISME 13:2306–2318

16. Ritcher T, Hazen T, Lam D, Coles J, Seidman J, You Y, Silbergeld E, Fraser C, Rasko D. 2018. Temporal variability of *Escherichia coli* diversity in the gastrointestinal tracts of Tanzanian children with and without exposure to antibiotics. mSphere 3:e00558–18 doi:101128/msphere00558-18

17. Fang X, Monk JM, Nurk S, Akseshina M, Zhu Q, Gemmell C, Gianetto-Hill C, Leung N, Szubin R, Sanders J, Beck PL, Li W, Sandborn WJ, Gray-Owen SD, Knight R, Allen-Vercoe E, Palsson BO, Smarr L. 2018. Metagenomics-based strain-level analysis of *Escherichia coli* from a time-series of microbiome samples from a Crohn’s disease patient. Front Microbiol 9:2559 doi:103389/fmicb 201802559

18. Chen DW, Garud NR. Rapid evolution and strain turnover in the infant gut microbiome. BioRxiv [preprint]. doi: https://doi.org/10.1101/2021.09.26.461856

19. Brito I, Yilmaz S, Huang K, Xu L, Jupiter S, Jenkins A, Naisilisili W, Tamminen M, Smillie C, Wortman J, Birren B, Xavier R, Blainey P, Singh A, Gevers D, Alm E. 2016. Mobile genes in the human microbiome are structured from global to individual scales. Nature 535:435–439 doi:101038/nature18927

20. Nicolas M, Bertrand X, Madec J. 2014. *Escherichia coli* ST131, an intriguing clonal group. Clin Microbiol Rev 27:543–574 doi:101128/cmr00125-13

21. Yamaji R, Rubin J, Thys E, Friedman C, Riley L. 2018. Persistent pandemic lineages of uropathogenic *Escherichia coli* in a college community from 1999 to 2017. J Clin Microbiol 56:e01834–17 doi:101128/jcm01834-17

22. Stoesser N, Sheppard AE, Pankhurst L, De Maio N, Moore CE, Sebra R, Turner P, Anson LW, Kasarskis A, Batty EM, Kos V, Wilson DJ, Phetsouvanh R, Wyllie D, Sokurenko E, Manges AR, Johnson TJ, Price LB, Peto TE, Johnson JR, Didelot X, Walker AS, Crook DW; Modernizing Medical Microbiology Informatics Group (MMMIG). Evolutionary history of the global emergence of the *Escherichia coli* epidemic clone ST131. mBio. 2016 Mar 22;7(2):e02162. doi: 10.1128/mBio.02162-15.

23. Codjoe FS, Donkor ES. 2017. Carbapenem Resistance: A Review. Med Sci (Basel).6(1):1. doi: 10.3390/medsci6010001.

24. Nobrega DB, Tang KL, Caffrey NP, De Buck J, Cork SC, Ronksley PE, Polachek AJ, Ganshorn H, Sharma N, Kastelic JP, Kellner JD, Ghali WA, Barkema HW. Prevalence of antimicrobial resistance genes and its association with restricted antimicrobial use in food-producing animals: a systematic review and meta analysis. J Antimicrob Chemother. 2021 Feb 11;76(3):561–575. doi: 10.1093/jac/dkaa443. PMID: 33146719.

25. Richter T, Michalski J, Zanetti L, Tennant S, Chen W, Rasko D. 2018. Responses of the human gut *Escherichia coli* population to pathogen and antibiotic disturbances. mSystems 3:e00047–18

26. Selzer P, Marhöfer R, Koch .2018. Applied Bioinformatics 2nd Ed Springer Cham: Switzerland

27. Tallarida, R.J. and Murray, R.B., 1987. Chi-square test. In Manual of pharmacologic calculations (pp. 140–142). Springer, New York, NY

28. Kurowski, K.M., Marusinec, R., Amato, H.K., Saraiva-Garcia, C., Loayza, F., Salinas, L., Trueba, G. and Graham, J.P., 2021. Social and environmental determinants of community-acquired antimicrobial-resistant *Escherichia coli* in children living in semirural communities of Quito, Ecuador. The American journal of tropical medicine and hygiene, 105(3), p.600.

29. Kozlov A. M., Darriba D., Flouri T., Morel B., Stamatakis A. (2019). RAxML-NG: a fast, scalable and user-friendly tool for maximum likelihood phylogenetic inference. Bioinformatics 35: 4453–4455. 10.1093/bioinformatics/btz305

